# Coconut inflorescence sap mediated synthesis of silver nanoparticles and its diverse antimicrobial properties

**DOI:** 10.1101/775940

**Authors:** M.K. Rajesh, K.S. Muralikrishna, Swapna S. Nair, B. Krishna Kumar, T.M. Subrahmanya, K.P. Sonu, K. Subaharan, H. Sweta, T.S. Keshava Prasad, Neeli Chandran, K.B. Hebbar, Anitha Karun

## Abstract

Green synthesis of nanoparticles (NPs) involves the use of diverse extracts of biological origin as substrates to synthesize nanoparticles and can overcome the hazards associated with chemical methods. Coconut inflorescence sap, which is unfermented phloem sap obtained by tapping of coconut inflorescence, is a rich source of sugars and secondary metabolites. In this study, coconut inflorescence sap was used to synthesize silver nanoparticles (AgNPs). We have initially undertaken metabolomic profiling of coconut inflorescence sap from West Coast Tall cultivar to delineate its individual components. Secondary metabolites constituted the major portion of the inflorescence sap along with sugars, lipids and, peptides. The concentration of silver nitrate, inflorescence sap and incubation temperature for synthesis of AgNPs were optimized. Incubating the reaction mixture at 40°C was found to enhance AgNP synthesis. The AgNPs synthesized were characterized using UV-Visible spectrophotometry, X-Ray Diffraction (XRD), Fourier Transform Infrared spectroscopy (FTIR), Field Emission Scanning Electron Microscopy (FESEM) and Transmission Electron Microscopy (TEM). Antimicrobial property of AgNP was tested in tissue culture of arecanut (*Areca catechu* L.) where bacterial contamination (*Bacillus pumilus*) was a frequent occurrence. Significant reduction in the contamination was observed when plantlets were treated with aqueous solutions of 0.01, 0.02 and 0.03% of AgNPs for one hour. Notably, treatment with AgNPs did not affect growth and development of the arecanut plantlets. Cytotoxicity of AgNPs was quantified in HeLa cells. Viability (%) of HeLa cells declined significantly at 10 ppm concentration of AgNP and complete mortality was observed at 60 ppm. Antimicrobial properties of AgNPs synthesized from inflorescence sap were also evaluated and confirmed in human pathogenic bacteria viz., *Salmonella* sp., *Vibrio parahaemolyticus,* and *Escherichia coli.* The study concludes that unfermented inflorescence sap, with above neutral pH, serves as an excellent reducing agent to synthesize AgNPs from Ag^+^.

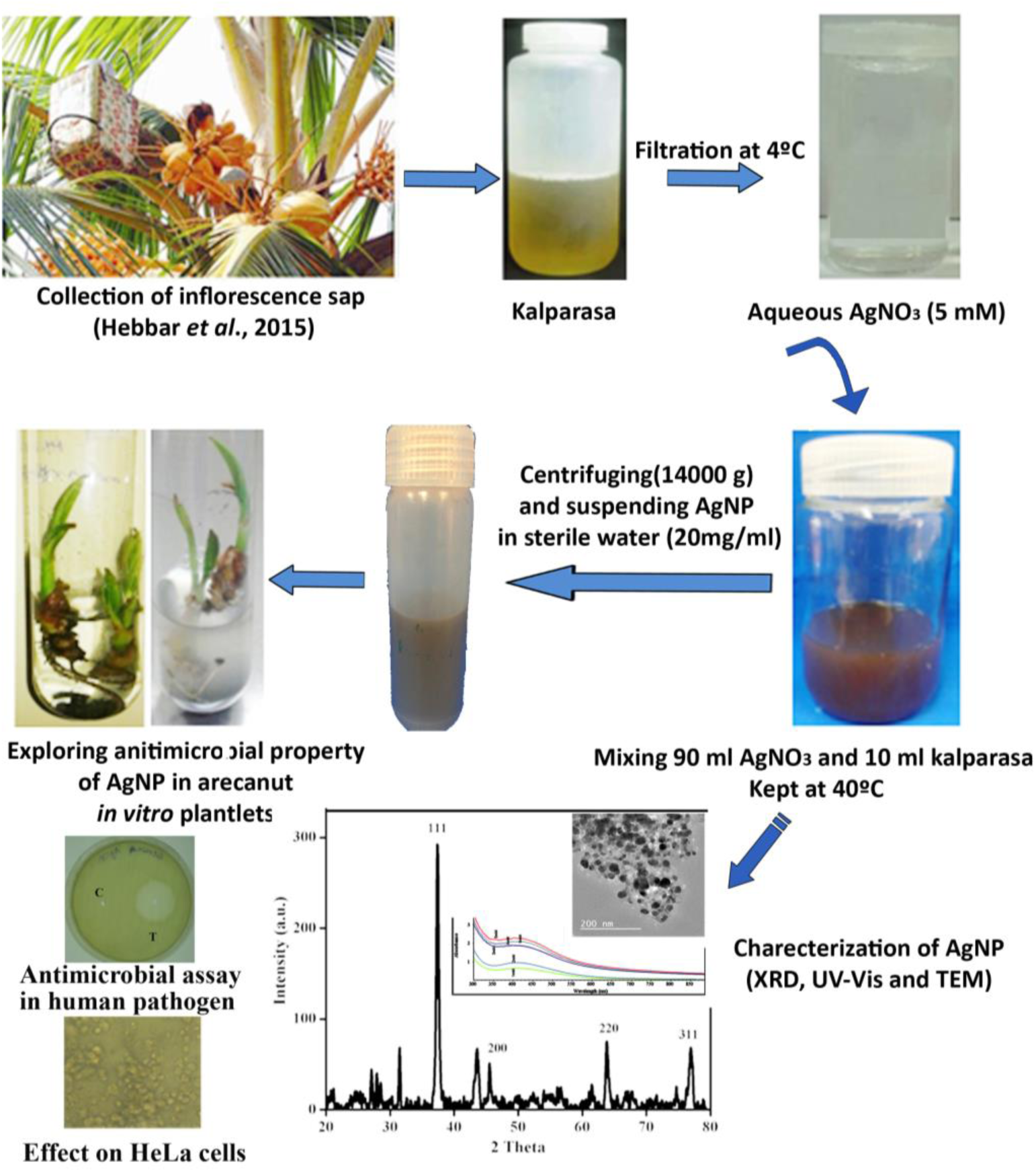
Graphical abstract

## 1. Introduction

In the recent years, syntheses of various nanoparticles (NPs), which range in size from 1-100 nm, have incited enhanced interest in the field of nanosciences as these NPs possess unique catalytic, optical, electronic, biological and magnetic properties. These nanomaterials find a repertoire of applications in different domains such as biosensors, electronics, catalysts, wastewater treatment, cosmetics, drug delivery, packaging and health (Li et al., 2011). Synthesis of NPs by conventional chemical or physical methods has limitations, particularly due to the environmental hazards associated with the process, the resources required and the time taken for the process (Gajbhiye et al., 2009; Shankar et al., 2004). Rapid, economical and eco-friendly techniques have thus evolved as alternative methods to chemical methods and to meet the growing requirement of NPs. Green synthesis or bioinspired synthesis of NPs, where metallic NPs are synthesized using sources of biological origin, is one such method. Extracts from plants and microorganisms, such as bacteria and fungi, have been used as substrates to synthesize NPs (Makarov et al., 2014; Singh et al., 2016; Abdelghany et al., 2018). However, culturing time and cross-contamination issues have restricted the application of bacteria and fungi in nanoparticle synthesis, to a limited extent. Plant mediated green synthesis has a great advantage over other biological methods as the synthesis is user-friendly and the synthesis is rapid (Ahmed et al., 2015). Plant-based substrates used for synthesis of NPs include extracts from leaves (Ahmed et al., 2016; Banerjee et al., 2014), fruits (Jain et al., 2009), seeds (Baghizadeh et al., 2015; Bar et al., 2009), peels (Abdelmonem and Amin, 2014), latex (Bar et al., 2009) and roots (Suresh et al., 2014). Plant extracts have been reported to serve both as reducing and capping agents during the synthesis of NPs (Bar et al., 2009). The availability of plant substrates in huge quantities makes plant extract mediated nanoparticle synthesis an apt alternative to the conventional methods (Zhang et al., 2016). The precise chemistry of green synthesis of NPs is still unclear given that plants comprise of a large number of compounds. Nonetheless, the reducing ability of plant extracts has been attributed to phytochemicals such as polyphenols (for e.g. catechines and stilbenes; Rodríguez-León et al., 2013), proteins (Firdhouse and Lalitha, 2013), amino acid residues (Awwadet al., 2013) and antioxidants (Ahmed and Sharma, 2012).

Extensive work has been carried out to evaluate antimicrobial properties of silver nanoparticles (AgNPs) with many reports have suggesting its superior antibacterial (Banerjee et al., 2014) and antifungal (Gajbhiyeet al., 2009) activities. The antimicrobial activity is highly dependent on the specific surface area of the NPs (Gupta and Silver, 1998) and a shape-dependent interaction of AgNPs with microbes has been established (Pal et al., 2007). Toxic effects of AgNPs has been primarily attributed to the penetration of microbial cell by silver ions released from AgNP resulting in a cascade of events such as membrane damage, DNA damage and reduced protein expressions (Hsueh et al., 2015).

Inflorescence sap or *neera*, a healthy drink consumed largely by the rural village folk, is tapped from coconut inflorescence (‘spadix’). A technique has been developed to collect inflorescence sap in its pure, hygienic and natural form using ‘coco sap chillers’ (Hebbar *et al*., 2015). Briefly, a ‘coco sap chiller’ makes use of ice cubes inside a specialized box which maintains the temperature (4 to 6°C) for 10-12 hrs to protect the collected inflorescence sap from getting fermented and also to maintain the sap free of foreign materials. Since it is a sap enrooted from the phloem, it is rich in sugars and other secondary metabolites. Coconut inflorescence sap has been reported to consist of reducing agents such as amino acids (0.25 g), proteins (0.16 g), phenolics (5.1 mg) and antioxidant activity (0.321 mM TE) per 100 ml (Augustine and Hebbar, 2014), which can be exploited for green routed synthesis of AgNPs.

Fresh inflorescence sap has not been tested previously for its efficacy to synthesize NPs and discovering such a novel biological source is advantageous. With this background, the present study was designed to green synthesize AgNPs using fresh coconut inflorescence sap, right after tapping. Apart from quantifying the basic parameters such as pH and sugars, the metabolomic profiling of coconut inflorescence sap was undertaken to delineate the individual components. Impacts of silver nitrate (AgNO3) concentration, the concentration of coconut inflorescence sap and incubation temperature were evaluated to optimize AgNP synthesis. Antimicrobial properties of AgNP particles synthesized were tested on arecanut (*Areca catechu* L.) *in vitro* cultures and human pathogenic bacteria and cytotoxic assessment were also carried out in HeLa cells.

## 2. Materials and Methods

### 2.1. Collection and processing of coconut inflorescence sap

The inflorescence sap (25-year-old West Coast Tall palm), which was collected in a coco-sap chiller under cool temperature as described by Hebbar et al. (2015), was brought to the laboratory in autoclaved glass bottles in an icebox. The sap was filtered using Whatmann No.1 filter paper inside a refrigerator (4°C). Fresh sap collected using this method comprise of total phenols (21.99 mg g-1), total flavonoids (0.96 mg g^-1^), free amino acids (901 mg g^-1^), vitamins such as ascorbic acid (13.45 mg g^-1^), niacin (14.86 mg g^-1^) and tocopherol (7.9 mg g^-1^) (Hebbar et al., 2018). In the freshly collected filtered sap, pH (inoLab WTW digital pH meter), reducing sugars (Nelson-Somogyi, 1952) and total sugar content (DuBois et al., 1956) were measured. The inflorescence sap was stored in refrigerator (4°C) for further analysis.

### 2.2. Metabolomic profiling of inflorescence sap

Metabolomic profiles were analyzed using freshly tapped coconut inflorescence sap. To the 50 µl of inflorescence sap, 10 volumes of 2:1 of chloroform: methane solution was added and vortexed for 10 min followed by sonication for 2 min. To this, 0.2 equivalent weight of water was added and centrifuged at 12000 rpm at 4°C for 5 min. Differentiated phases were collected, dried using the speed vacuum and the content was re-suspended in 100 µl of 0.1% formic acid water. An aliquot of the sample (1:10 diluted) was injected in the column of LC-MS (Agilent). The LC had 50 mm x 2.1mm, 1.8-micron Zorbax C18 Eclipse RRHD Plus column with a flow rate of 0.2 mL min^-1^ and a run time of 40 minutes. The MS had independent positive and negative modes with a mass range from 50-1000 and functions based on IDA-EPI (independent data acquisition-enhanced product ion) method.

### 2.3. Optimization of AgNO_3_ and inflorescence sap concentrations for the synthesis of nanoparticles

Normally, green synthesis of NPs follows a procedure wherein the aqueous plant extract is mixed with aqueous solution of the suitable metal salt and incubating for specific time for the completion of reduction process. In this study, the filtered inflorescence sap was directly used as reducing agent to synthesize AgNPs using AgNO_3_ (SIGMA) as precursor. The reaction mixture (100 ml) comprised of 5 mM aqueous AgNO_3_ (90 ml) and coconut inflorescence sap (10 ml). The reaction mixture was prepared in sterile glass bottles (Borosil) covered with lid and kept in different conditions *viz*., room temperature (RT, 27±2°C), refrigerator (4°C) and incubator (40°C). Incubation time was 24 hrs in all the treatments. Synthesis of silver nanoparticle was confirmed by a color change of the solution from colorless to reddish-brown.

Different concentrations of aqueous AgNO_3_ (1.0, 2.0, 3.0, 4.0, 5.0 and 6.0 mM) were prepared in sterile water. Reaction mixtures prepared in a glass bottle with a tight lid comprised of inflorescence sap (2.5, 5.0, 10.0 and 20.0 ml) and AgNO_3_ (97.5, 95.0, 90.0 and 80.0 ml) in a total volume of 100 ml. The reaction mixture was kept at the selected temperature in an incubator for the complete synthesis of NPs for a duration of 24 hours.

### 2.4. Characterization of AgNP particles

#### 2.4.1. UV-visible spectrometer

UV-Vis spectrum of the resulting solution was observed by scanning in a UV-Vis Spectrophotometer (Thermo Fisher Scientific) between 900 −300 nm with a resolution of 1 nm. Based on the UV spectrophotometric observations, the best responding combination was identified and used for further characterization of NPs.

#### 2.4.2. Fourier Transform Infrared (FT-IR) Spectroscopy

FT-IR spectra ranging from 600 cm^-1^ to 4000 cm^-1^ of AgNPs were recorded by FT-IR (Jasco FT/IR-6300, Tokyo, Japan) spectrometer for identifying biomolecules /phytochemicals present in the inflorescence sap. An aliquot of the purified sample of AgNPs was crushed with well dried KBr and pressed into pellets for recording the spectra.

#### 2.4.3. Field Emission Scanning Electron Microscopy (FESEM)

The reaction mixture was centrifuged at 14,000 rpm and pellets were washed thrice with sterile water. The centrifugation was repeated followed by ethanol washing and dried. A minute amount of AgNP was well dispersed in ethanol by sonication for FESEM (Nova-Nano SEM 600 FEI, Netherlands) observation. This solution was used for the characterization of NPs using FESEM analysis.

#### 2.4.4. Transmission Electron Microscopy (TEM)

Morphology and size distribution of AgNP were studied by TEM observations. A small fraction of AgNP was suspended thoroughly in ethanol using sonicator for 30 min. A drop of the suspension was then placed on a copper grid and dried thoroughly. The grid loaded with the sample was observed under TEM (JEOL, JEM3010, JAPAN). Well dispersed areas were selected and images were captured to study the structure and size distribution.

#### 2.4.5. X-ray Diffraction (XRD) analysis

Study of crystalline nature of AgNP was performed by using powder x-ray diffraction (XRD; Bruker, D8 advance). The required amount of sample was placed onto a sample holder and scanned for 1 hour between 2θ range from 10° to 90°.

### 2.5. Assessment of anti-microbial properties of AgNPs

#### 2.5.1. Antimicrobial bioassay of AgNPs in arecanut(Areca catechu L.) in vitro plantlets

The AgNPs synthesized was suspended in sterile water at 20 mg ml^-1^ concentration after centrifuging and washing with sterile water and used directly to treat the arecanut *in vitro* plantlets, derived from immature inflorescence cultures (Karun et al., 2004). Frequent bacterial contamination was encountered during the regeneration process. The identification of bacteria was carried out using 16S rRNA sequencing. Plantlets in contaminated liquid medium were washed in sterile water and kept in AgNP solutions of varying concentration (0.01, 0.02 and 0.03%) for 60 minutes. Later, plantlets were thoroughly washed in sterile water for three to four times and subsequently transferred to fresh liquid nutrient medium (Y3 medium with 30 g L^-1^ sucrose, 2 mg L^-1^ each of BAP and NAA and 1 g L^-1^ activated charcoal). Cultures were kept under observation for six days.

#### 2.5.2. Bactericidal activity of AgNPs

Stock cultures of standard bacterial isolates (*Salmonella enterica* ATCC 14028; *Escherichia coli* ATCC 25922; *Vibrio parahaemolyticus* AQ4037) maintained in −80°C in a deep freezer at NITTE University Centre for Science Education and Research, Mangalore, India was used to study the bactericidal activity of the AgNPs by well diffusion method. These bacterial isolates were aerobically grown in Luria-Bertani medium (HiMedia, India) at 37°C on a rotary shaker at 200 rpm. A lawn of each isolate was prepared uniformly by spreading bacterial culture on individual Muller-Hinton agar (HiMedia, India) and 6 mm diameter wells were made using gel puncture. Ten microliters of different concentration of AgNPs (0, 1, 2.5, 10 and 25 ppm) were added into their respective wells and incubated at 37 °C for 24 hours. After incubation, the obtained zone of inhibition (ZOI) was measured.

#### 2.5.3. In vitro cytotoxicity studies of AgNPs in HeLa cells

HeLa cells were cultured in Dulbecco’s Modified Eagle’s Medium (DMEM) (Gibco, Thermo Fischer Scientific) supplemented with 10% (v/v) Fetal Bovine Serum (FBS), antibiotic antimycotic solution (100X) (1 ml 100 mL^-1^; Sigma), gentamicin (40 mg mL^-1^; Sigma) 40 µl 100 mL^-1^ and 2 mM glutamine (Sigma, USA). Cells were allowed to grow to form a monolayer in a 37°C incubator (Panasonic, Japan) supplemented with 5% CO_2_. For estimation of the cell viability against AgNP, HeLa cells were seeded at 3 X10^3^cells per well in 96-well plates and pre-cultured for two days. HeLa cells were treated with a series of AgNPs ranging from 0 ppm to 100 ppm along with cell control and incubated for 12 h in a 37°C incubator supplemented with 5% CO_2_. After treatment, HeLa cells were washed twice with 1 X PBS to remove the floating cells. Then, 200 µL MTT reagent was added to each of the wells and incubated for another 3 hrs under the same conditions. After the removal of the medium, 200 µl of DMSO was added and incubated for 5 minutes under shaking to dissolve formazan crystals and absorbance measured at 570 nm. Three independent experiments were conducted in duplicates to maintain the accuracy and average values are depicted as a graphical representation.

## 3. Results and discussion

### 3.1. Metabolomic profiling of coconut inflorescence sap

Freshly collected and filtered sap had a pH of 7.26 and it consisted of 14.18% of total sugars and 0.62% reducing sugars. The obtained values are well within the range (Hebbar et al., 2018).

In order to obtain a deeper understanding of the mechanism of green synthesis of AgNPs, we have initially undertaken untargeted metabolomic profiling of fresh coconut inflorescence sap. Secondary metabolites, sugars, lipids, and peptides were detected in fresh inflorescence sap. The MS profile revealed that in the positive mode, fresh coconut inflorescence sap had 64 % of secondary metabolites, 9% sugars, 12% lipids/fats and 9% peptides, whereas in the negative mode it was 33, 20, 11 and 9%, respectively (**Fig. 1**).

**Fig. 1.**
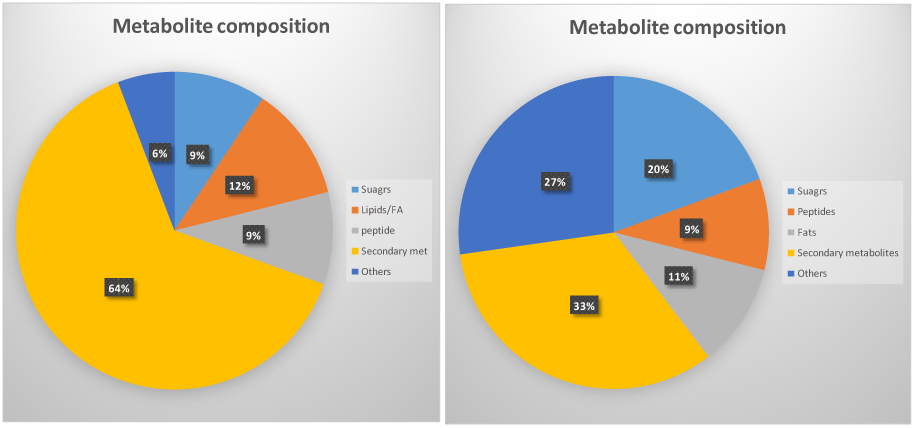
Anionic and cationic metabolites in the coconut inflorescence sap

The major metabolites identified were polyphenols of the class flavonoids such as 3,3’,4’,5,6,8-hexamethoxyflavone (22 ppm), xanthones including gentisin (22 ppm), alkaloids such as (R)-boschniakine (20 ppm), atherosperminine (18 ppm), 4-coumaroyl-2-hydroxyputrescine (18 ppm) and dipeptides like glutamylmethionine (10 ppm). An extensive assortment of molecules, encompassing numerous low molecular weight compounds to high molecular weight proteins have been implicated to assay a key role in the green synthesis of NPs (Marslin et al., 2018). These molecules include amino acids (Awwad et al., 2013), polyphenols (Rodríguez-León et al., 2013; Hebbar et al., 2018), organic acids (Ramteke et al., 2013), sugars (Shankar et al., 2004), terpenoids (Kaviya et al., 2011), alcoholic compounds (Liu et al., 2018), alkaloids (Velmurugan et al., 2014), antioxidants (Ahmed and Sharma, 2012) and proteins (Firdhouse and Lalitha, 2013). It has been postulated that these molecules could be involved both in the reduction of metal ions into NPs and also assist their subsequent stability (Makarov et al., 2014). The compound which has been commonly reported to partake in plant-based green synthesis of NPs are polyphenols, especially flavonoids (Marslin et al., 2018). Given the predominance of polyphenols like flavonoids and xanthones in fresh coconut inflorescence sap, it could be assumed that these compounds could be mainly involved in synthesis of NPs. Many studies have reported the role of polyphenols in the promotion of green synthesis of metal NPs (Tao et al., 2016; Jigaysa and Rajput, 2018; Singh et al., 2019; Latif et al., 2019).

### 3.2. Optimization of concentration of AgNO_3_, inflorescence sap volume and synthesis condition

Suitable conditions to synthesize NPs were initially standardized wherein the reaction mixture (90 ml of 5mM AgNO_3_ and 10 ml of coconut inflorescence sap) was kept in refrigerator (4°C), room temperature (27±2°C) and in incubator (40°C). In all the conditions, addition of inflorescence sap to the AgNO_3_ solution led to the appearance of brown color which indicates the formation of AgNPs. However, the absorbance in 400-425 nm range after 24 hrs was highest for the samples kept in an incubator at 40 °C as compared to those kept at refrigerator or in room temperature (**Fig. 2**).

**Fig. 2.**
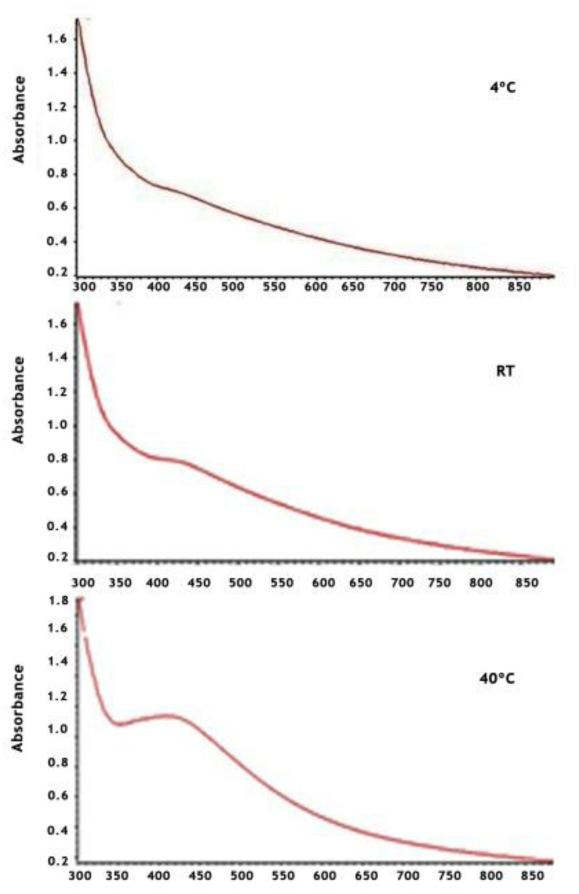
The UV-visible spectrum of AgNP synthesized using fresh inflorescence sap of coconut under different conditions *viz*., room temperature (RT), refrigerator (4°C) and incubator (40°C) with a synthesis time of 24 hrs.

In another set of experiments, the reaction mixture was formed with varying concentrations of AgNO_3_ (1, 2, 3, 4, 5 and 6 mM) and volumes of coconut inflorescence sap (2.5, 5, 10 and 20 ml in a reaction mixture of 100 ml). This translates the inflorescence sap concentration to 2.5, 5, 10 and 20%, respectively in each treatment. As the concentration of AgNO_3_ increased, the absorbance at 415 nm also increased and increment was highest with 5 mM of AgNO_3_. In general, absorbance of the reaction mixture at 415 nm increased as the volume of inflorescence sap increased and maximum absorbance was observed in reaction mixture with 90% of 5 mM AgNP and 10 % of inflorescence sap as indicated by dark colour (**Fig. 3**). The experiment indicates that 48 mg of Ag+ present in 90 ml of 5 mM AgNO_3_ could be reduced with the addition of 10 ml of inflorescence sap.

**Fig. 3.**
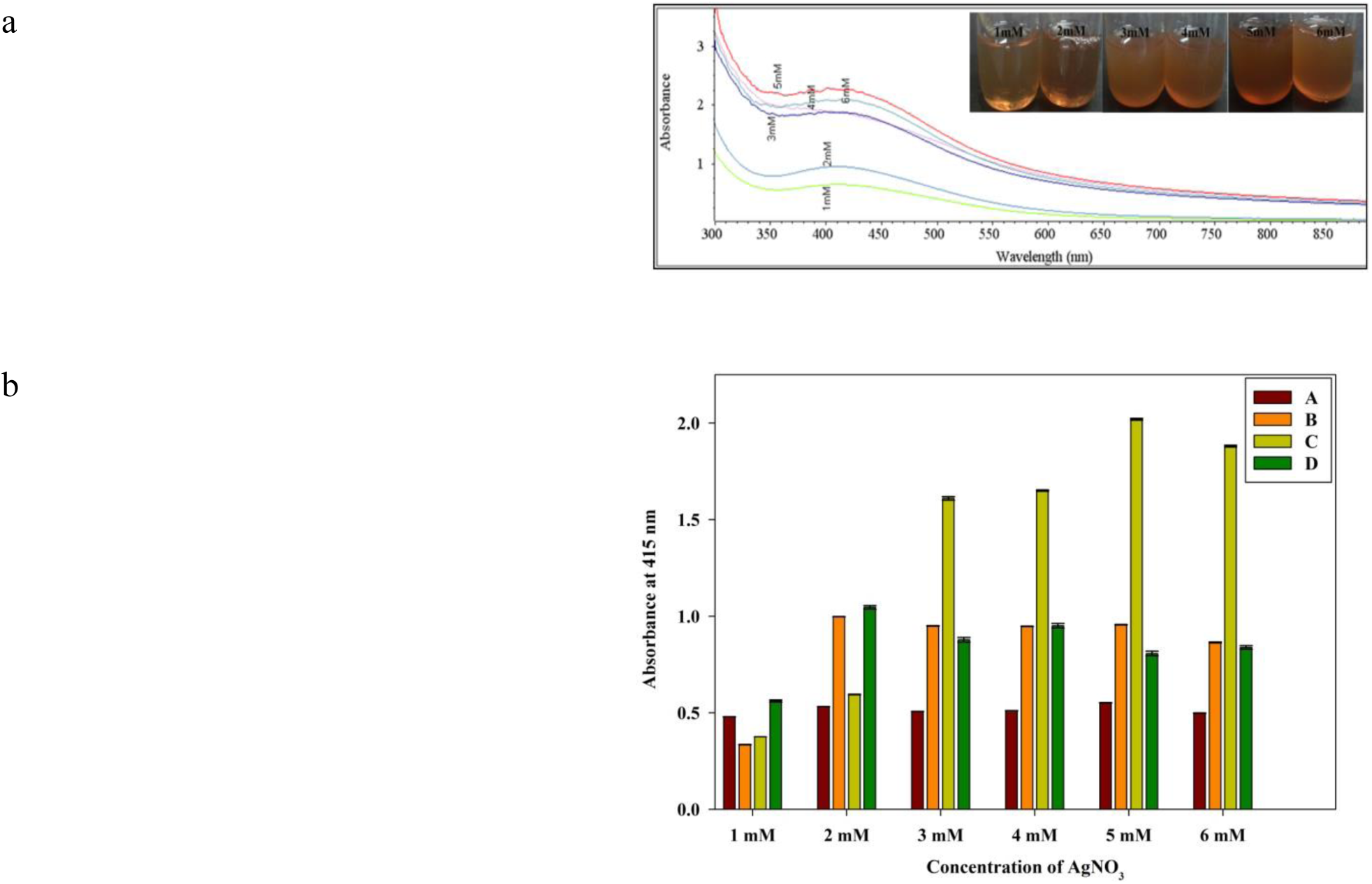
The UV-visible spectrum of AgNP synthesized using fresh inflorescence sap of coconut. (a) UV-visible spectra from 300 to 900 nm with a resolution of 1 nm of the reaction mixture (100 ml) comprising of 90 ml of varying the concentrations of aqueous AgNO_3_ (1, 2, 3, 4, 5 and 6 mM) and 10 ml of inflorescence sap. (b) Average absorbance of the reaction mixture with varying concentrations of AgNO_3_ and inflorescence sap. Reaction mixture ‘A’ consisted of 97.5 ml of AgNO_3_ (1, 2, 3, 4, 5 and 6 mM) and 2.5 ml of inflorescence sap, ‘B’ with 95 ml and 5 ml, ‘C’ with 90 ml and 10 ml while ‘D’ with 80 ml and 20 ml, respectively. The reddish-brown colour indicates the formation of AgNP (inset photo).

The change of the color of the reaction mixture to brownish red has been attributed to the excitation of surface Plasmon resonance, a characteristic feature of AgNP (Banerjee et al., 2014; Veeraswamy et al., 2011). Increase in incubation temperature has been reported to enhance AgNP production (Chahardooli et al., 2014), which corroborates our results since synthesis of AgNPs was more in 40°C compared to 4°C or room temperature. As the reaction temperature increases, the rate of reaction increases and secondary reduction process stops as most of the Ag+ engaged in the formation of nuclei and lead to small-sized nanoparticles (Song and Kim, 2009).

The change in color and absorbance signifies the size of the particles in the reaction mixture (Tripathy et al., 2010). The experiment indicates that 48 mg of metallic silver present in 90 ml of 5 mM AgNO_3_ could be reduced with the addition of 10 ml of inflorescence sap. A fresh inflorescence sap consists 0.62 g of reducing sugar (Table 1) and 0.245 g of amino acids, 0.16 g of protein, 5.1 mg of phenolics and 0.321 mM TE of antioxidant activity per 100 ml (Augustine and Hebbar, 2014). This translates to ∼62 mg of reducing sugars, 24.5 mg of amino acids, 510 µg of phenolics and antioxidant activity of 0.032 mM TE, which might have played their part in the reduction process. This phenomenon can be attributed to the increasing rate of spontaneous nucleation significantly accelerating the growth rate of silver nanoparticles (Abdel-Mohsen et al., 2012), reflecting a higher amount of AgNPs prepared due to the increased amount of reducing biomolecules available at a higher dosage. Other sources from coconut like liquid endosperm (Elumalai et al., 2014), shell (Sinsinwar et al., 2018) and oil (Govarthanan et al., 2016) were used effectively to synthesize silver nanoparticles. Similar results were reported by Sathishkumar et al. (2009) and Dubey et al. (2010). Properties and stability of nanoparticles, in general decided by the size of the nanoparticles (Wu et al., 2014). Above neutral pH in the reaction mixture leads to the formation of nanoparticles with reduced size and with more stability (**He et al., 2017**). In the present study; reducing agent, inflorescence sap in its pure form has above neutral pH, thus adding a spontaneous advantage in nanoparticle synthesis.

**Table 1.** Chemical properties of fresh inflorescence sap of coconut.

| Parameter | Value |
| --- | --- |
| pH | 7.26±0.2 |
| Reducing sugars (g 100 ml<sup>-1</sup>) | 0.62±0.1 |
| Total sugars (g 100 ml<sup>-1</sup>) | 14.18±0.5 |

Values are represented as a mean of three replications.

### 3.3. Characterization of AgNP

Characterization of AgNPs was undertaken carried out in NPs resulting from a reaction mixture comprised of 90 ml of 5 mM AgNO_3_ and 10 ml of inflorescence sap as the response was better compared to rest of the combinations as indicated through UV-Vis spectrophotometer analysis.

#### 3.3.1. Fourier Transform Infrared (FT-IR) Spectroscopy

The functional groups present in the inflorescence sap responsible for the reduction of Ag+ were identified by FT-IR analysis. The FT-IR spectra of AgNPs indicated the presence of clear peaks at 3329 cm^-1^, 2932 cm^-1^, 1621 cm^-1^, 1516 cm^-1^, 1393 cm^-1^ and 1086 cm^-1^ (Fig. 4). The obtained peaks corresponded to OH group of alcohols or phenols, C-H of alkanes, C≡C group of alkynes, C=C group of alkenes, N-O group, C=O, C-N group of aliphatic amines, C-O group of alcohols, carboxylic acids, esters or ethers. These groups correspond to the presence of phenols, flavonoids, amino acids, proteins, vitamins as the unfermented sap is rich source of these metabolites (Hebbar et al., 2018; Borse et al., 2007; Xia et al., 2011) These groups have been reported to be involved in suppression of Fenton’s reaction, which is driven by superoxides and is a chief source of reactive oxygen species (ROS), thus catalysing metallic NP formation (Marslin et al., 2018).

**Fig. 4.**
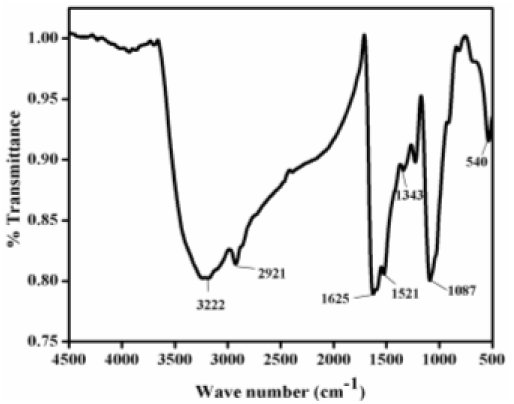
FTIR analysis of AgNPs

The FT-IR spectroscopy analysis in the present study confirms the role of biomolecules present in coconut inflorescence sap as a reducing agent in the synthesis of AgNP. Peaks depict stretching vibrations responsible for biomolecules such as polyphenols and other natural compounds, such as fatty acids and proteins, which can act as both reducing and stabilizing agents to form AgNPs (Huang *et al*., 2007; Shankar *et al*., 2004). The ability of these natural metabolites to serve as capping agents offer further advantage. In the present study, the untargeted metabolome profiling revealed the presence of natural compounds like flavonoids, xanthones, and alkaloids in coconut inflorescence sap providing further proxy to the hypothesis that these natural compounds act as reducing, stabilizing and capping agents in the synthesis of AgNPs. Accordingly, the presence of elevated quantities of reducing/stabilizing agents prevents NP aggregation thus resulting in smaller NPs (Marslin et al., 2018).

#### 3.3.2. Field Emission Scanning Electron Microscopy (FESEM)

The morphology of colloidal AgNP powder was characterized by FESEM analysis. Majority of the AgNP produced using inflorescence sap were spherical in shape as revealed from FESEM images (Fig. 5). This is also well evidenced by the peaks obtained in the FTIR analysis of AgNPs.

**Fig. 5.**
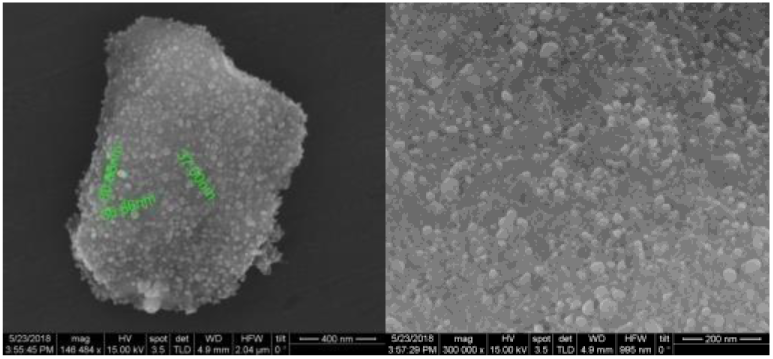
FESEM images of AgNPs

#### 3.3.3. Transmission Electron Microscopy (TEM)

The nature of structure and size distribution of AgNP was examined by TEM studies. Figure 6 reveals the predominantly spherical shaped AgNP. The bulk of the particles was spherical with smooth (thin) shell and poly-dispersed with the particle diameter ranging from 10 nm to 30 nm. The TEM image provided an indication of agglomerates of small grains and a few dispersed NPs, corroborating the results acquired by FESEM (Guzmán et al., 2009). Dark shades observed on the surface of the NPs could be attributed to the organic compounds present in the coconut inflorescence sap which, in addition to primarily serving as a reducing agent in the synthesis of NPs, could also serve as a capping agent, preventing the aggregation of NPs (Rai et al., 2012).

**Fig. 6.**
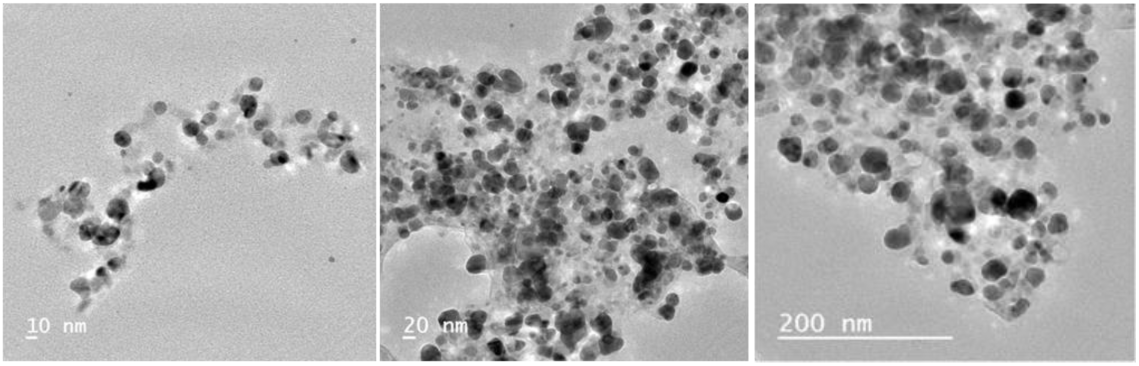
TEM images of AgNPs

#### 3.3.4. X-ray Diffraction (XRD) analysis

Crystalline nature of the AgNP particles was revealed in XRD pattern. The peak at 37.5° for (111), 45.5° for (200), 63.8° for (220), and 77° for (311) planes could be attributed to the planes of face-centered-cubic crystals of AgNP (Fig. 7). Distinct peaks revealed in the XRD pattern at 2*Ѳ* values gives ample evidence for the presence of face-centered cubic lattice in AgNP crystals (de Jesus Ruiz et al., 2017; Kumar et al., 2015). Other minor peaks were also observed, which could be associated with the biomolecules present on the surface of the AgNP (Kumar et al., 2017), which could be accountable for the reduction of Ag^+^ and stabilization of resultant nanoparticles (Ibrahim, 2015)

**Fig.7.**
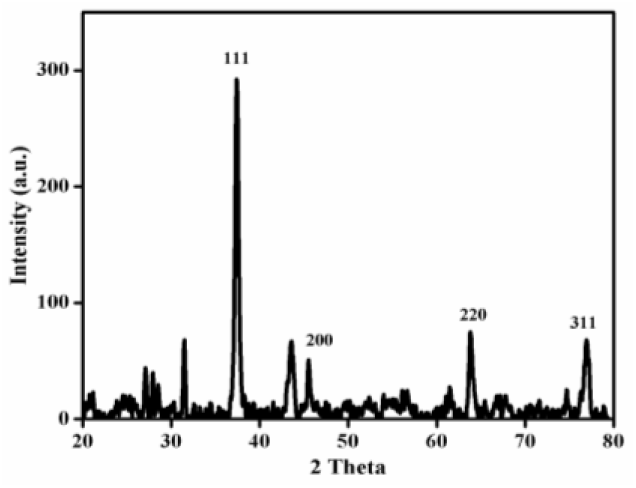
X-ray diffraction (XRD) pattern of synthesized AgNPs

### 3.4. Efficacy of AgNP against bacterial contamination in arecanut in vitro cultures

The antibacterial property of the AgNP particles was tested on *in vitro* plantlets of arecanut derived from immature inflorescence culture in our laboratory. Frequent bacterial contamination of tissue cultured plantlets was identified as *Bacillus pumilus* (GenBank accession number MN339666). Contaminated plantlets were treated with different concentrations of AgNP solution (Fig. 9). Bacteria did not recur six days post-treatment when plantlets treated in AgNP solution of concentration as low as 0.01% for one hour and subsequent transfer into the fresh culture medium (Fig. 9). Results indicate that contamination could be regulated without any visible effect on growth and development of the plantlet.

**Fig. 9.**
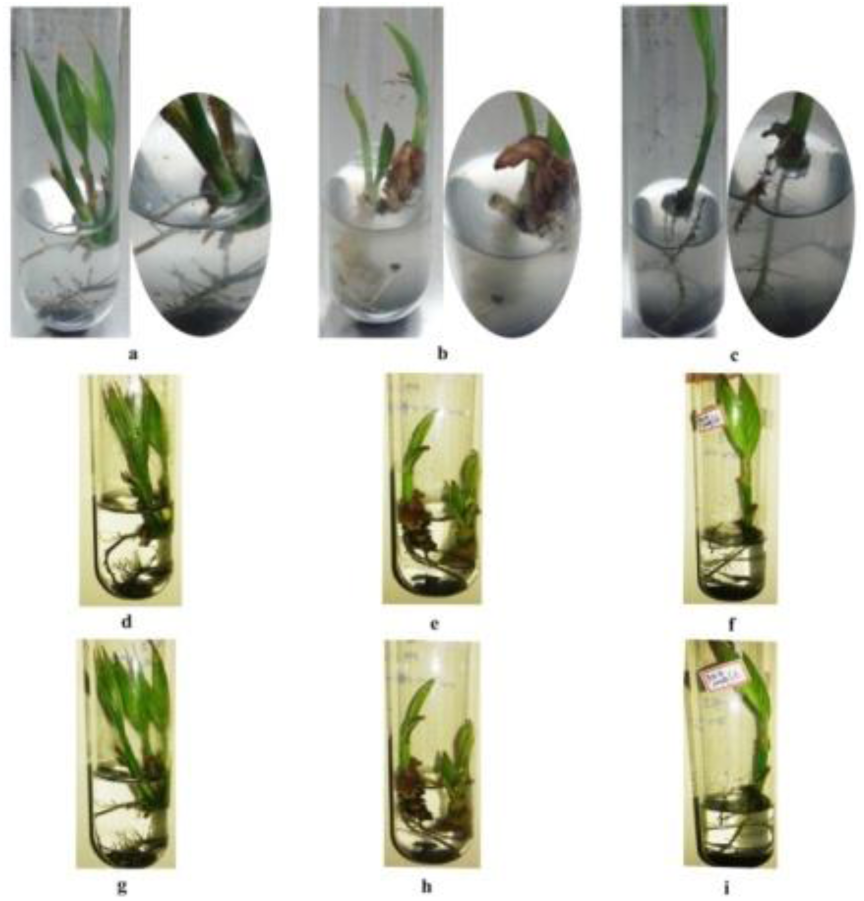
Bacterial contamination observed in arecanut (*Areca catechu* L.) plantlets and its control using green synthesized AgNP. Contaminated plantlets were washed in sterile water and then kept in aqueous solution consist of 100 (a), 200 (b) and 300 mg/l (c) of AgNP for 60 minutes. Plantlets then transferred to liquid medium (Eeuwens Y3) with growth regulators (BAP and NAA) and observed for three (d, e and f) and six days (g, h and i) post-treatment.

Contamination is a major obstacle for *in vitro* culturing of plants and arecanut (*Areca catechu* L.) is no different. This makes it essential to find ways out to eliminate bacterial contamination since it poses a significant challenge in the final outcome. *B. pumilus* is a Gram-positive bacteria and reported to be a contaminant in tissue cultured plants (Rossini and Standardi, 1990). In the present study, the recovery of contaminated arecanut plantlets could be achieved with the use of AgNP indicating its antimicrobial property. Use of AgNPs, in the form of soaking of explants or incorporation into the tissue culture medium, has been reported to eliminate microbial contamination during *in vitro* culturing of olive (Rostami and Shahsavar, 2009), almond x peach hybrid rootstocks (Arab et al., 2014), Norfolk-Island pine (Sarmast et al., 2011) and rubber (Moradpour et al., 2016).

### 3.5. Efficacy of AgNP against select pathogenic bacteria

Results of the bactericidal assay carried out using coconut inflorescence mediated synthesized AgNPs on human pathogenic bacteria such as *Escherichia coli, Salmonella sp.* and *Vibrio parahaemolyticus* AQ4037 are represented in Table 2. *In vitro* assay, where the zone of inhibition (mm) was measured in the presence of different concentrations of AgNPs, revealed that concentration of 10 ppm and above had the highest effect in three of the tested organisms (Fig. 10), while the lower level of 1 ppm had no impact on any of the cultures.

**Fig. 10.**
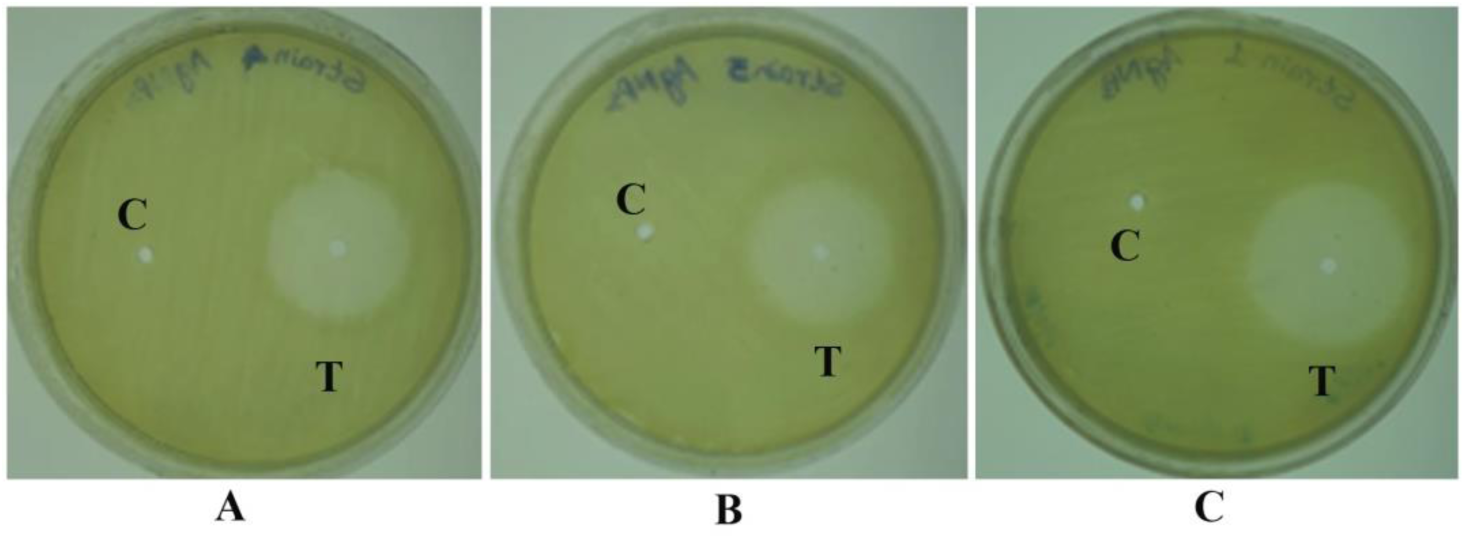
Effect of AgNPs on three different pathogenic microorganisms at 10 ppm concentration (A) *Salmonella* (B) *Vibrio parahaemolyticus* AQ4037 (C) *Escherichia coli*. Zone of inhibition was measured as millimeter ± standard deviation for three independent experiments.

**Table 2.** Effect of AgNPs on pathogenic microorganisms. Zone of inhibition was measured as millimeter ± standard deviation for three independent experiments

| Test organism | Zone of inhibition (mm) at different concentration (ppm) |  |  |  |  |
| --- | --- | --- | --- | --- | --- |
|  | 0 | 1.0 | 2.5 | 10.0 | 25.0 |
| *Escherichia coli* | 0 | 0 | 15±1.2 | 25±1.54 | 25±.67 |
| *Salmonella sp.* | 0 | 0 | 9.5±.65 | 20±.93 | 20±.28 |
| *Vibrio parahaemolyticus* AQ4037 | 0 | 0 | 12±.84 | 20±.26 | 20±.1.1 |

The bactericidal property of the silver nanoparticles has been attributed to the release of tiny particles of silver which get attached to the bacterial cell wall, leading to protein denaturation and cell death (He et al., 2017).

### 3.6. In vitro cytotoxicity of AgNPs

The inflorescence sap mediated synthesized AgNP were subjected to cytotoxicity assay on HeLa cells. Results indicated significant dose-dependent anti-proliferative activity. Lower levels of concentrations viz., 1 and 5 ppm did not affect the viability of HeLa cells significantly, while viability (%) declined significantly at 10 ppm concentration and complete mortality was observed at 60 ppm and beyond till 100 ppm tested (Fig. 11). The results acquired from the present study can be corroborated with previous evidence for the cytotoxic effect of bioinspired AgNP synthesis against HeLa cancer cell lines viz., from leaf extracts of *Solanum muricatum* (Gorbe et al., 2016), *Punica granatum* (Sarkar and Kotteeswaran, 2018). It has been reported that nanoparticles have potential to selectivity bind and target cancer cells, making them special in the therapeutic field (Cho et al., 2008).

**Fig. 11.**
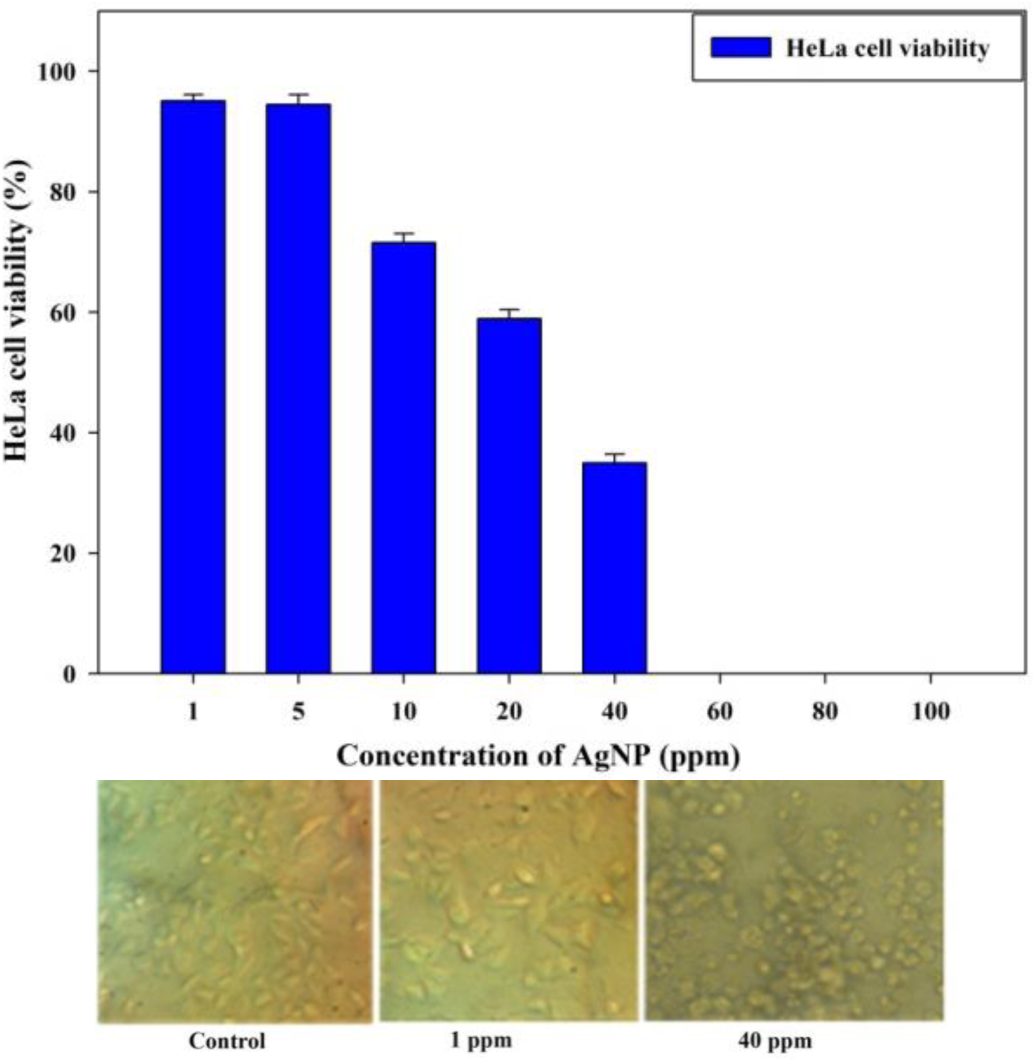
Effect of biosynthesized AgNP on viability (%) of HeLa cells.

## 4. Conclusions

The study highlights the efficiency of fresh inflorescence sap from coconut, in its pure and hygienic form, to synthesize AgNPs without addition of external stabilizers or reducing agents. Preparation of the substrate for the green synthesis was much simpler as compared to other plant extracts which usually require processes such as drying, powdering, and solvent extraction. Thus the inflorescence sap tapped from coconut inflorescence is a ‘ready to use’ source to synthesize AgNPs through much ‘greener’ process as it bypasses the solvent extraction process or addition of external stabilizers or capping agent. Similar to the antimicrobial properties of AgNP widely reported earlier, inflorescence sap mediated AgNP showed efficient antimicrobial activities and were used effectively to reduce the bacterial contaminations in arecanut *in vitro* plantlets.

## Abbreviations

AgNPs: Silver nanoparticles BAP: Benzyl Amino Purine
FESEM: Field emission scanning electron microscope
FTIR: Fourier transformation infrared
MIC: Minimum inhibitory concentration NAA: Naphthalene Acetic Acid
NPs: Nanoparticles XRD: X-ray diffraction

## Acknowledgments

Authors are grateful to Indian Council of Agricultural Research (ICAR) for funding this research.

